# Graph Drawing-based Dimensionality Reduction to Identify Hidden Communities in Single-Cell Sequencing Spatial Representation

**DOI:** 10.1101/2020.05.05.078550

**Authors:** Alireza Khodadadi-Jamayran, Aristotelis Tsirigos

## Abstract

With the rapid growth of single cell sequencing technologies, finding cell communities with high accuracy has become crucial for large scale projects. Employing the current commonly used dimensionality reduction techniques such as tSNE and UMAP, it is often difficult to clearly distinguish cell communities in high dimensional space. Usually cell communities with similar origin and trajectories cluster so closely to each that their subtle but important differences do not become readily apparent. This creates a problem for clustering, as clustering is also performed on dimensionality reduction results. In order to identify such communities, scientists either perform broad clustering and then extract each cluster and perform re-clustering to identify sub-populations or they over-cluster the data and then merging the clusters with similar gene expressions. This is an incredibly cumbersome and time-consuming process. To solve this problem, we propose K-nearest-neighbor-based Network graph drawing Layout (KNetL, pronounced like ‘nettle’) for dimensionality reduction. In our method, we use force-directed graph drawing, whereby the attractive force (analogous to a spring force) and the repulsive force (analogous to an electrical force in atomic particles) between the cells are evaluated, and the cell communities are organized in a structural visualization. The coordinates of the force-compacted nodes are then extracted, and we employ dimensionality reduction methods, such as tSNE and UMAP to unpack the nodes. The final plot, a KNetL map, shows a visually-appealing and distinctive separation between cell communities. Our results show that KNetL maps bring significant resolution to visualizing and identifying otherwise hidden cell communities. All the algorithms are implemented in the iCellR package and available through the CRAN repository. Single (i) Cell R package (iCellR) provides great flexibility at every step of the analysis pipeline, including normalization, clustering, dimensionality reduction, interactive 2D and 3D visualizations, batch alignment or data integration, imputation, and interactive cell gating tools, which allow users to manually gate around the cells.

## INTRODUCTION

Oftentimes, using current commonly used dimensionality reduction methods for single cell analysis such as tSNE [1], UMAP [2], diffusion maps [3], and TriMap [4], it is rather difficult to increase the resolution of the dimensionality reduction to such an extent as to be able to visualize different cell communities with a high level of distinction. Lower resolution in both visualization and clustering means that many cell communities with subtle but important differences will either falsely resemble one another or remain hidden and unidentified. This can specifically be challenging for identifying cell-communities in large-scale, complex projects like Human Cell Atlas (HCA) [5]. To address this issue, investigators employ a rather cumbersome process: re-clustering the broadly clustered cell communities individually and manually browsing through the different clustering results. For instance, it is common to re-cluster T cells by first removing the non-T cell communities in order to be able to achieve a higher resolution and identify different T-cell sub-populations. However, this process is often error prone, since the broadly-clustered T-cells often include cells belonging to other communities due to low resolution.

To address this problem, we introduce a KNN-based Network graph drawing Layout (KNetL) for dimensionality reduction. KNetL map takes advantage of graph theory to present a structural visualization capable of representing cell communities with an amplified resolution. Dimensionality reduction plays a crucial role in the analysis of single-cell data, and therefore, a higher resolution is necessary to identify cell communities and define the properties of the cells. Our approach will have many applications from studying cancer heterogeneity to identifying novel cell communities and their functions.

## METHODS

KNN-based Network graph drawing Layout (KNetL) map finds K-Nearest Neighboring cells for every cell or node (root cell) and creates a link (edge) between the neighboring cells and the nodes by calculating their cell-cell distance to create a graph/network. This graph is then used to create a force-directed layout such as Fruchterman and Reingold layout algorithm [7]. The coordinates of the nodes in the layout are then extracted to perform dimensionality reduction using one of the common methods such as tSNE or UMAP to refine the position of the tightly compacted nodes. The force (resolution) of the system can be modified by adjusting the k values in KNN (Figure 1A-E).

**Figure 1.**
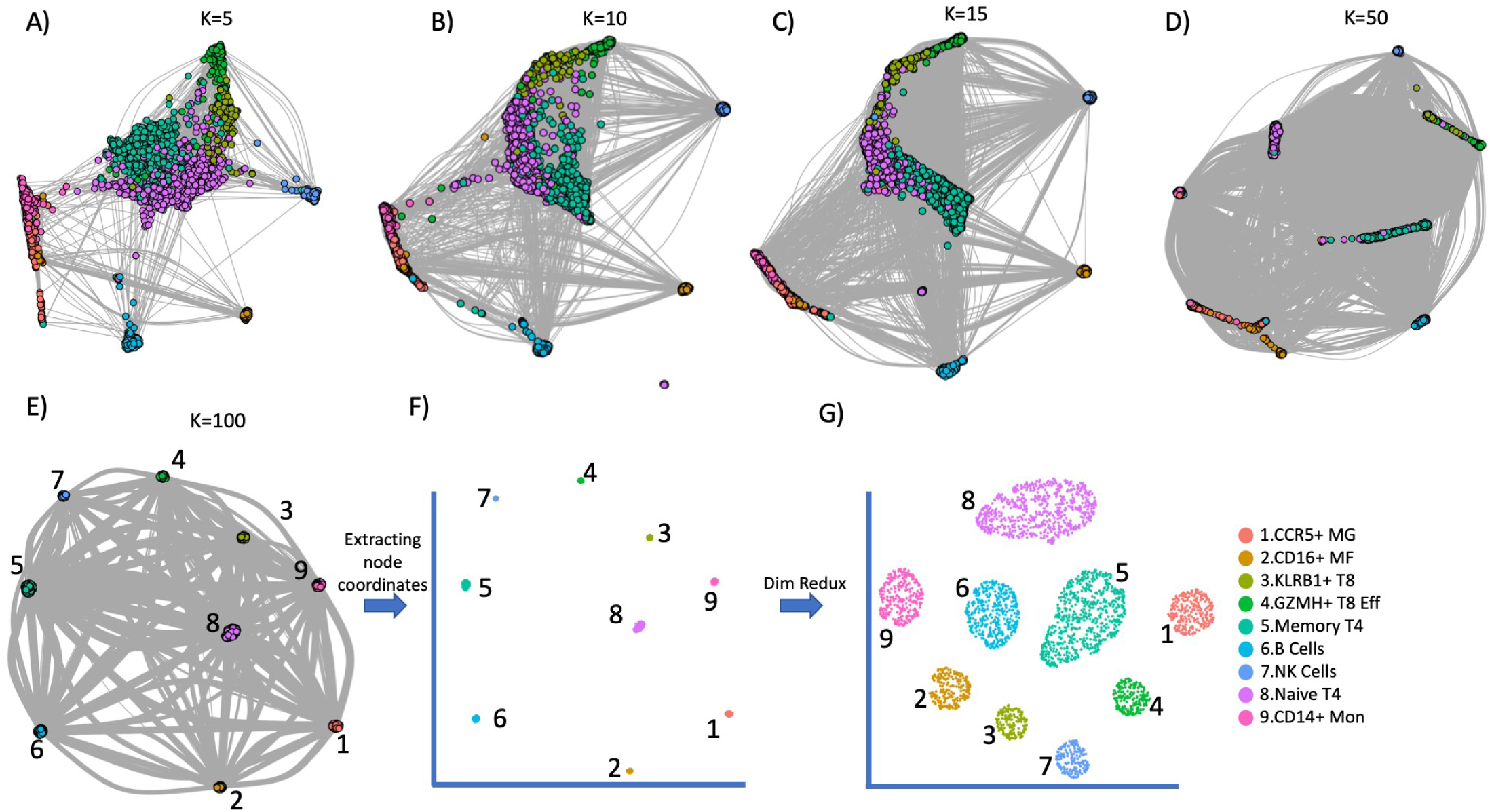
A) Raw KNetL plot of a sample PBMC data showing 9 clusters using K value of 5. Euclidian distance calculated on the first 20 PCs. B) Raw KNetL plot of a sample PBMC data showing 9 clusters using K value of 10. C) Raw KNetL plot of a sample PBMC data showing 9 clusters using K value of 15 D) Raw KNetL plot of a sample PBMC data showing 9 clusters using K value of 50. E) Raw KNetL plot of a sample PBMC data showing 9 clusters using K value of 100. Euclidian distance calculated on the first 20 PCs. Links are removed. F) Raw KNetL map coordinates. As shown in the figure all the cell communities are in distinct clusters but in tightly compacted space. G) KNetL map after unpacking the tightly compacted nodes using tSNE. As shown in the figure all the cell communities are in distinct clusters.

### Creating network edges

To create cell-cell links (edges) used in KNetL maps, we perform a Principle Component Analysis (PCA) on the expression matrix and then calculate the cell-cell distances using Euclidian distance. Then K Nearest Neighboring cells are found for every cell to create links. The KNN links can be created in two ways:

a. In a combined/joint fashion explained in Combined Coverage Correction Alignment (CCCA) and Combined Principle Component Alignment (CPCA) algorithms [6]. This could be useful for batch alignment using combined integration. In this method if K is equal to ten, ten links from every condition will be created for each cell. For instance, if there are two conditions there will be 20 links for very cell.
b. In a non-combined fashion explained in Coverage Correction (CC) algorithms [6]. In this method if K is equal to ten, it means ten links will be created for every cell regardless of the number of conditions.

The forces between the links are calculated by Euclidian distance (E value) gained from the first few principle components (20 by default).

### Graph Drawing

The links created using one of the techniques explained above are then used to make a graph. In this method, each cell is treated as an object (nodes/vertices) and the pairwise relations between the objects are obtained using the links (edges). This network can then be used for graph drawing using graph layout algorithms. The KNetL function implemented in iCellR allows users to choose from multiple different layout algorithms, but employs the force-directed Fruchterman and Reingold algorithm [7] by default. In force-directed algorithms the attractive force is analogous to the spring force based on Hooke’s law, while the repulsive force is analogous to the electrical force of atomic particles based on Coulomb’s law. The effect of these forces are meant to balance the energy of the system by moving the nodes [7].

### Dimensionality Reduction

Increasing the k values in KNN can adjust the resolution, however the nodes drawn using the force-directed graph drawings with high k values can also be tightly compacted. Lower k values don’t cause this issue, but they also often fail to capture details. To address this issue, we extract the coordinates of the cells (nodes) from the spatial representation created by graph drawing (Figure 1F) and perform a dimensionality reduction algorithm such as tSNE and UMAP to adjust the positioning of the nodes for a more refined visualization (Figure 1G).

### Adjusting for Resolution and Clustering Based on KNetL Maps

KNetL map is sensitive and must be adjusted in order to fully have grasp on the number of cell communities. Just like in microscopes, it is important to zoom in and out for an intended resolution in order to view the required amount of details. This can be done by adjusting the “k” parameter in the function. To zoom in decrease the k value; to zoom out increase the k value. Generally, KNetL is most effective with a k parameter somewhere between 100-600. The “dims” parameter also plays a role, and we tend to get better results when it’s set to 20. After adjusting the k value, in order to get a clear clustering comparable with this analysis, it is best to cluster the cell communities using the KNetL map coordinates rather than the principle components.

## RESULTS

To study the sensitivity of KNetL maps in distinguishing between different cell communities we used two data sets. A commonly analyzed sample PBMC data (a single sample) and a more complex PBMC data with nine samples from different sequencing technologies to show the KNetL map results on batch aligned data.

### KNetL Map Amplifies Distinction Between Cell Communities

To compare KNetL map to tSNE and UMAP, we used commonly analyzed data with known cell communities to visualize the localization of the cells in the plots. Our results show that KNetL map can significantly distinguish the different cell communities with clear separation (Figure 2). These communities can be further divided into subpopulations by adjusting the resolution (k and dims). To show the accuracy of the boundaries, we plotted the expression of some marker genes using KNetL plots (Figure 3), tSNE and UMAP plots (Figure S1 and S2). For instance, as shown in the Figure 3, clusters 8, 5, 3 and 4 are high CD3E (T Cells), clusters 3 and 4 are CD8+ and cluster 4 is high in GZMH. Clusters 5 and 8 are T4 cells, and cluster 5 has higher CCR7.

**Figure 2.**
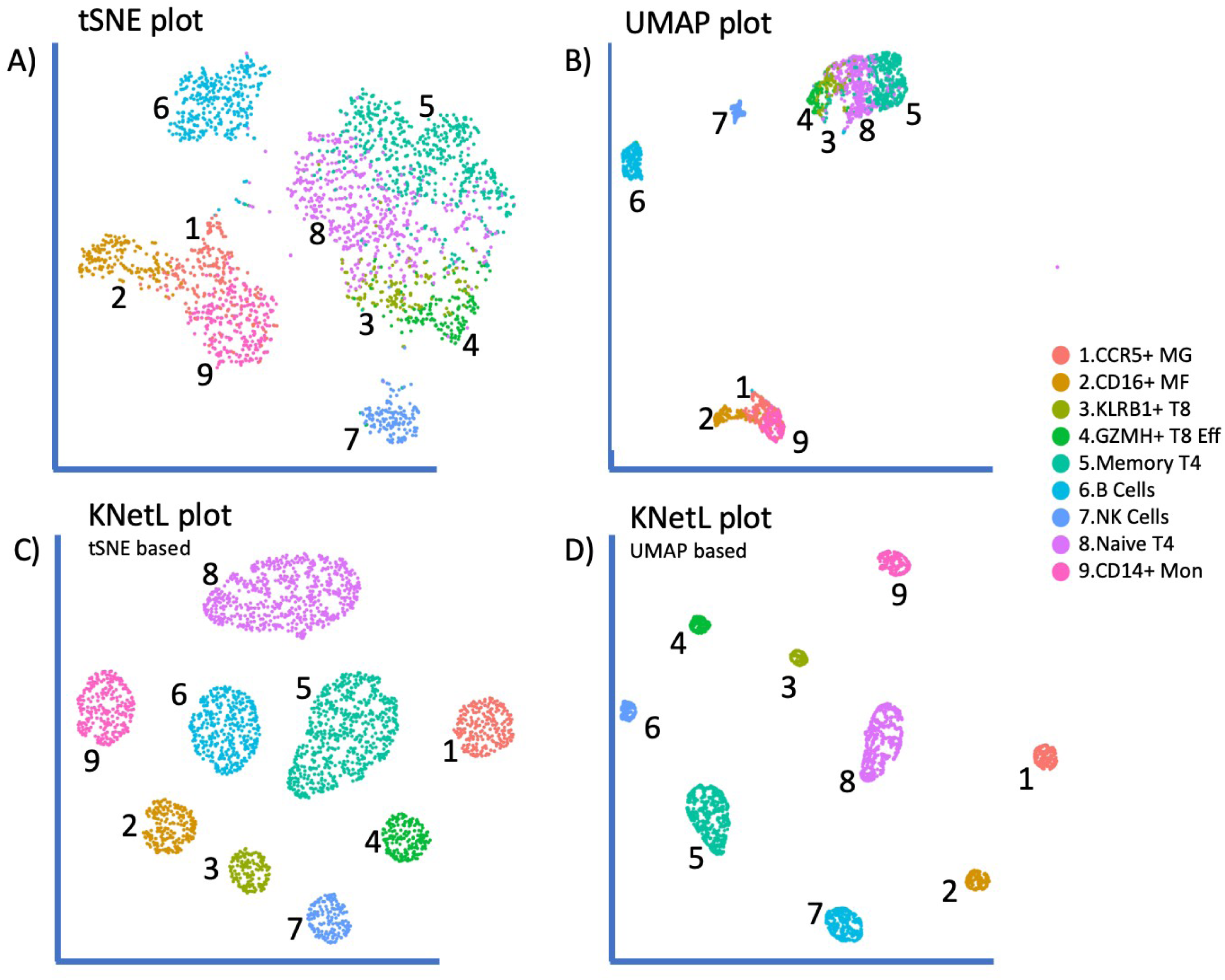
A) tSNE plot showing the 9 cell communities with less clear boundaries. B) UMAP plot showing the 9 cell communities with less clear boundaries. C) KNetL plot showing the 9 cell communities with clear boundaries. The nodes were unpacked using tSNE. D) KNetL plot showing the 9 cell communities with clear boundaries. The nodes were unpacked using UMAP.

**Figure 3.**
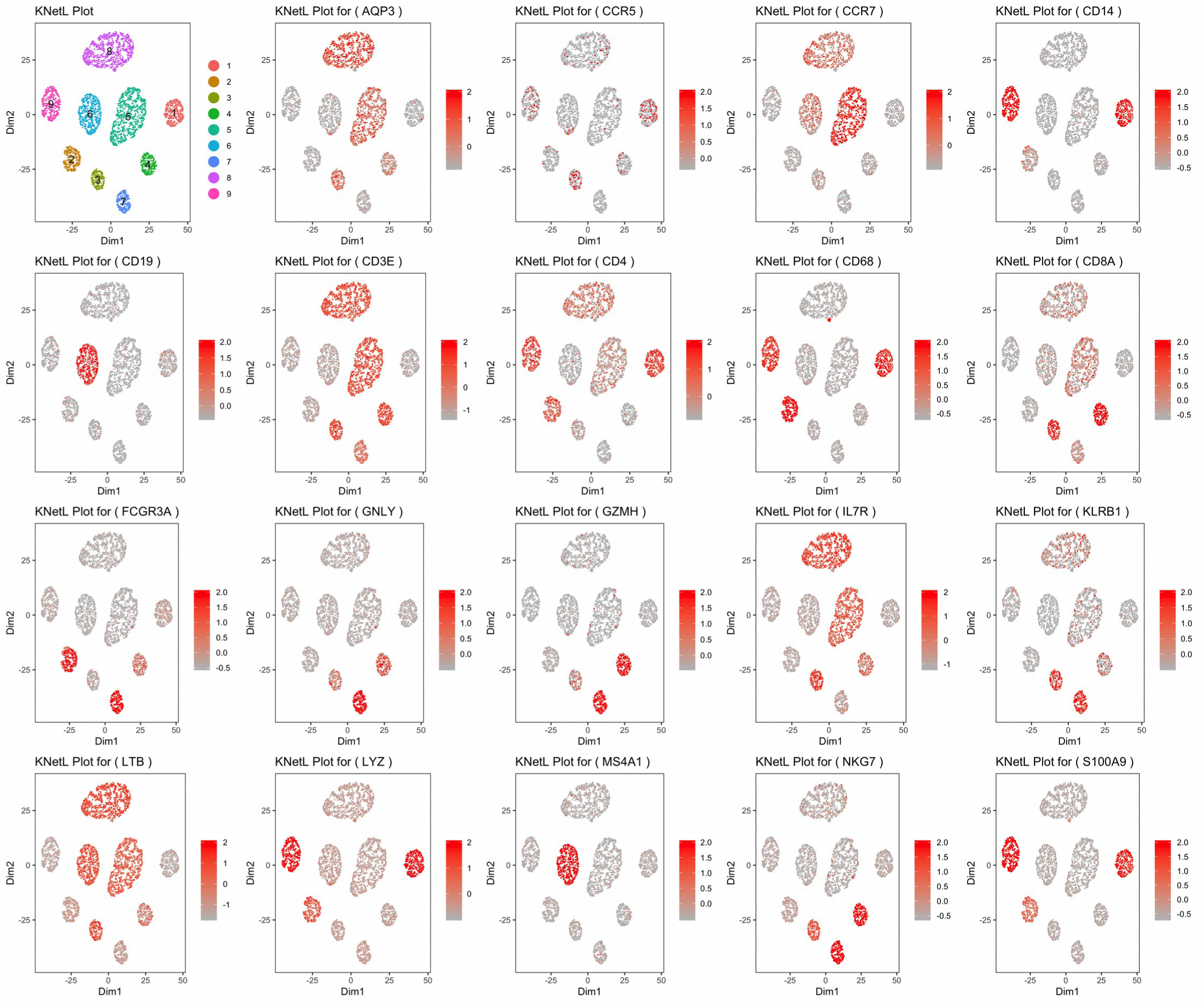
KNetL maps showing the accuracy of the boundaries by heatmapping the expression of some marker genes. As seen in the figure the cell communities are clearly distinguishable and easy to differentiate.

### KNetL Map Improves Batch Alignment Results

To test KNetL map on more complex data, we used 9 PBMC samples sequenced using different technologies (three 10x Chromium v2, 10x Chromium v3, CEL-Seq2, Drop-seq, inDrop, Seq-Well and SMART-Seq). These samples have been provided by the Broad Institute to study batch differences [8]. All the nine samples were then aligned or integrated using three different methods:

a. Combined Principal Component Alignment (CPCA) using iCellR [6].
b. Multiple Canonical Correlation Analysis (MultiCCA) using Seurat [9].
c. Mutual Nearest Neighbors (MNN) using scran [10].

The results were then plotted using UMAP and KNetL map plots shown in Figure 4 and the alignment of the nine samples using KNetL map is shown in Figure 5. Our results indicate that the cell communities are well distinguishable using KNetL plots. We have provided a heatmap of some marker genes in Figure 6.

**Figure 4.**
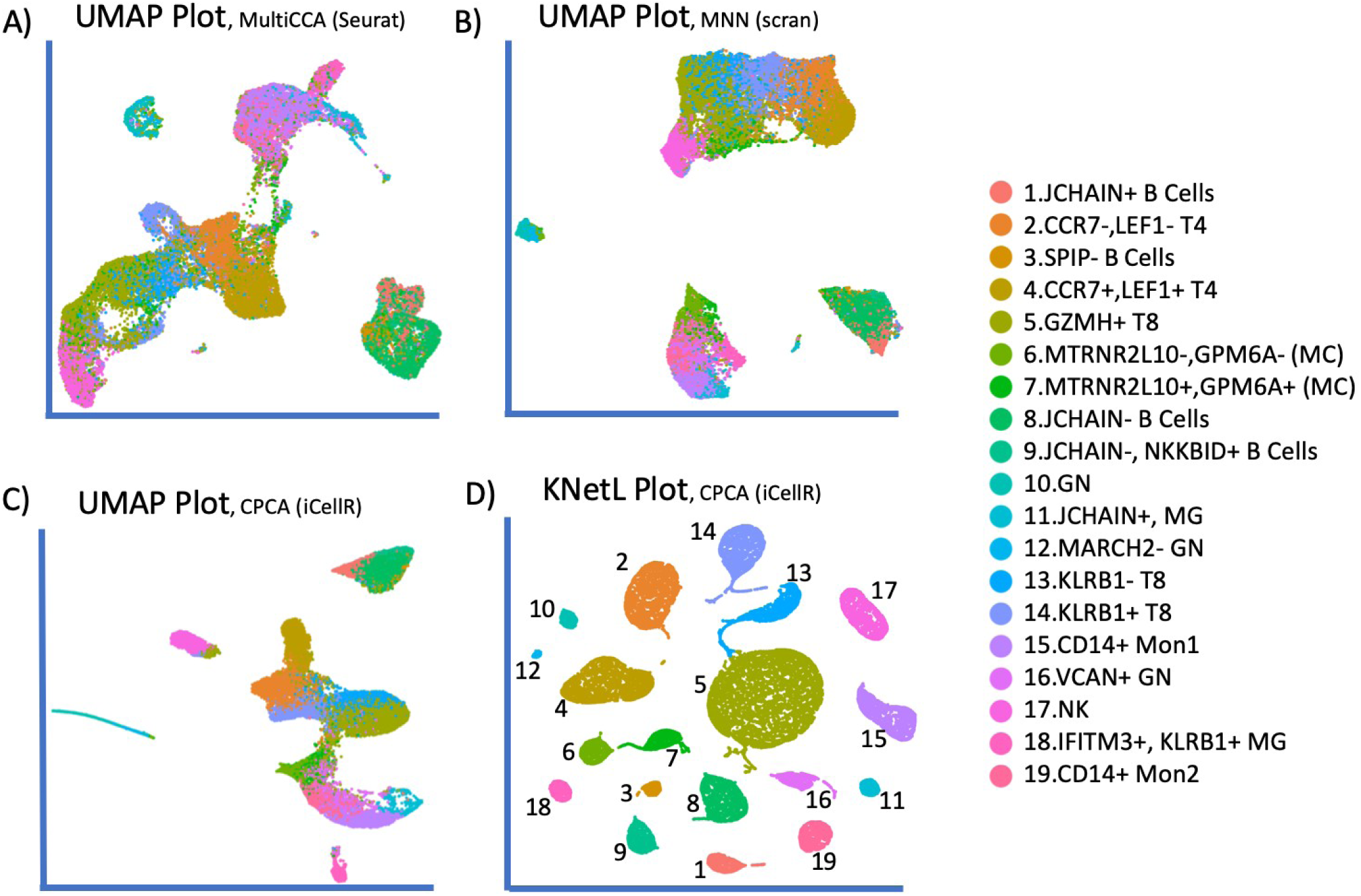
A) UMAP plot showing 19 cell communities from 9 PBMC samples batch aligned using MultiCCA. B) UMAP plot showing 19 cell communities from 9 PBMC samples batch aligned using MNN. C) UMAP plot showing 19 cell communities from 9 PBMC samples batch aligned using CPCA. D) KNetL plot showing 19 cell communities from 9 PBMC samples batch aligned using CPCA. The KNetL plot is UMAP based and with a k value of 400. As seen in the figure, all the cell communities have clear boundaries and easy to differentiate.

**Figure 5.**
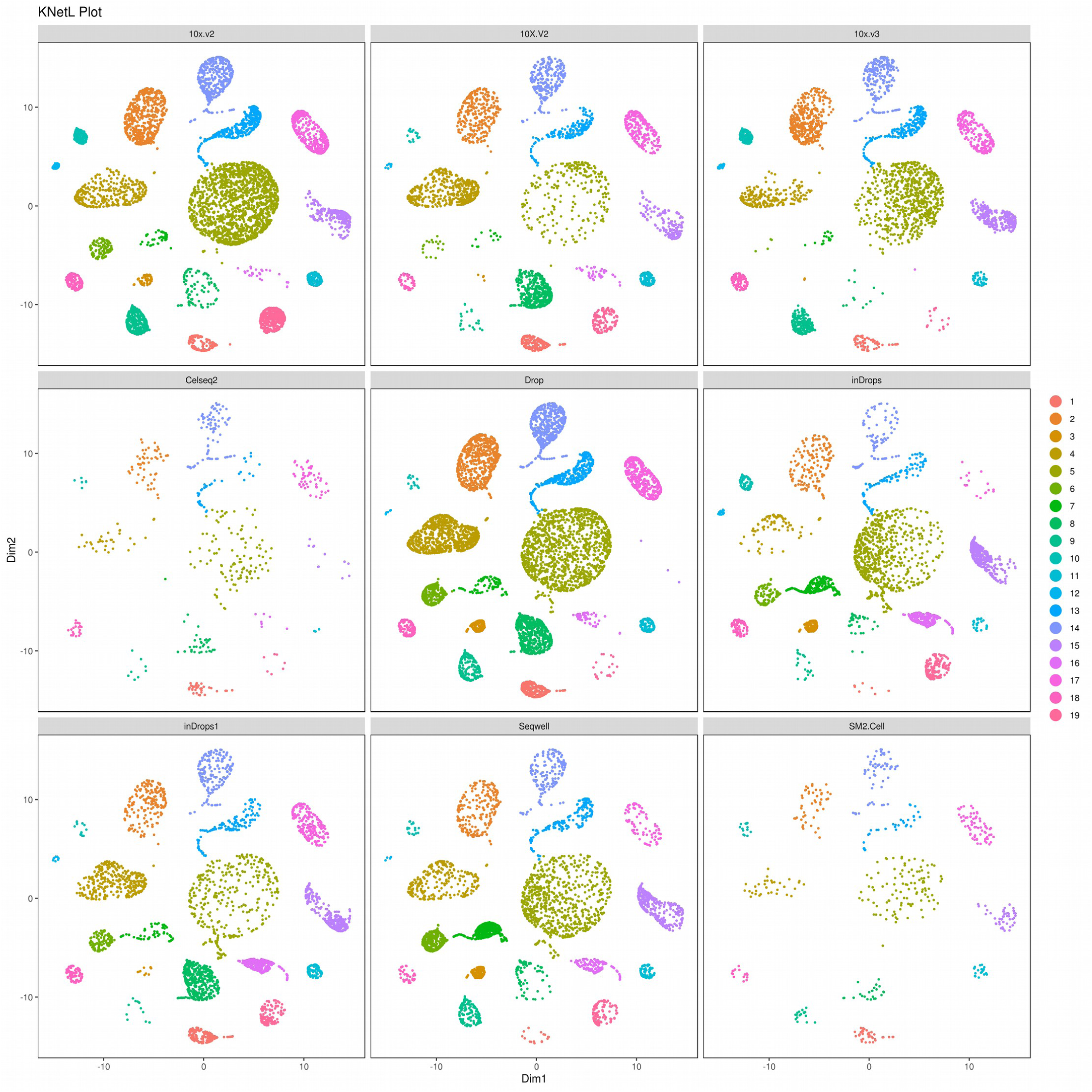
KNetL plot showing the alignment of the 9 PBMC samples. As seen in the figures the cell communities from all the 9 samples are clearly aligned.

**Figure 6.**
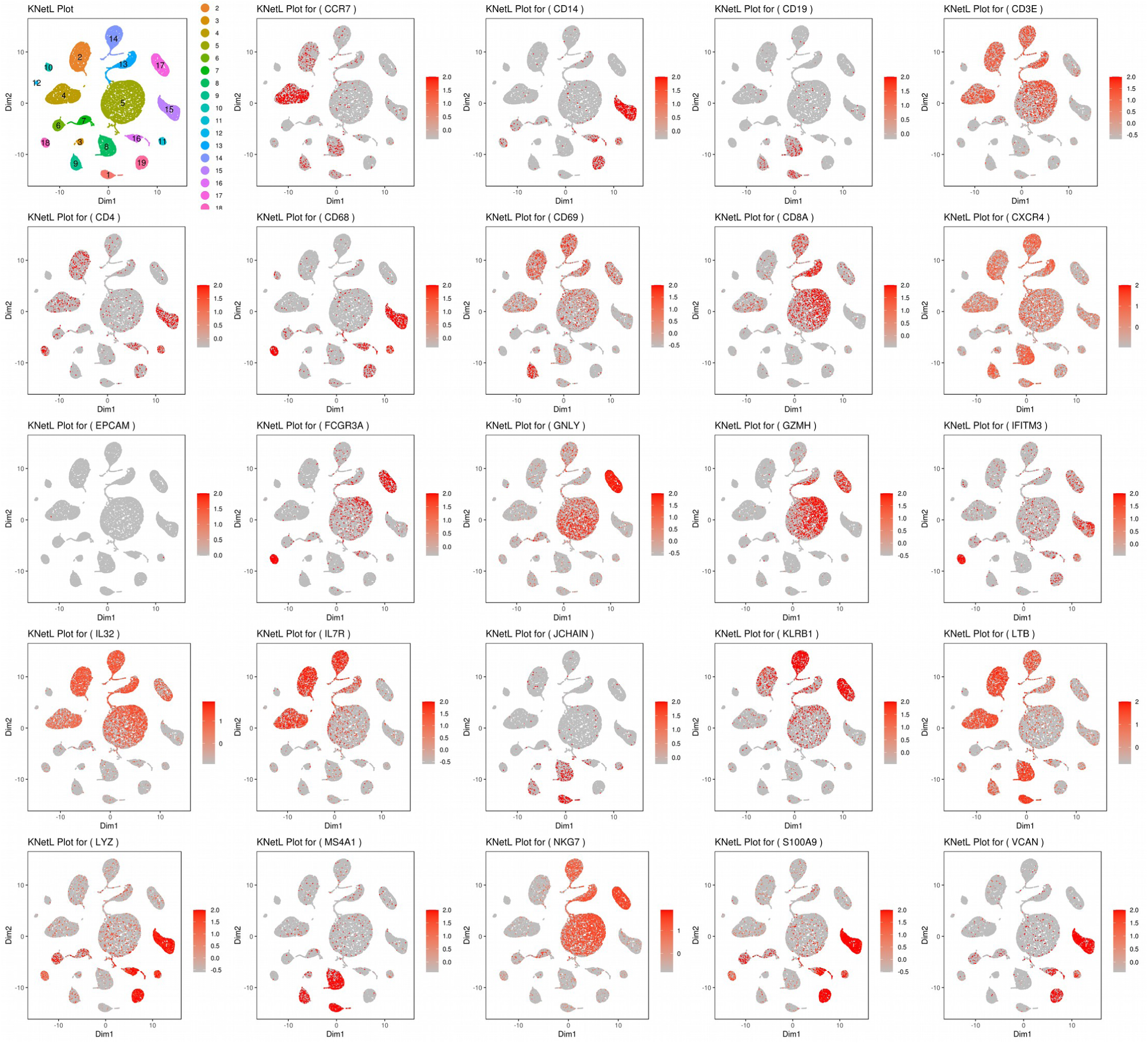
KNetL maps showing the accuracy of the boundaries by heatmapping the expression of some marker genes. As seen in the figure the cell communities are clearly distinguishable and easy to differentiate.

## DATA ACCESSION AND CODES

The PBMC data used for coverage correction can be downloaded from the 10x genomics website (https://support.10xgenomics.com/single-cell-gene-expression/datasets).

For connivance all these datasets can also be downloaded from here (https://genome.med.nyu.edu/results/external/iCellR/data/).

All the codes and algorithms are written in R and are available from iCellR package from CRAN (https://cran.r-project.org/package=iCellR).

In addition, the pipelines generating the figures in the results are in supplementary R scripts and the GitHub page for iCellR is (https://github.com/rezakj/iCellR).

---

**Figure S1.**
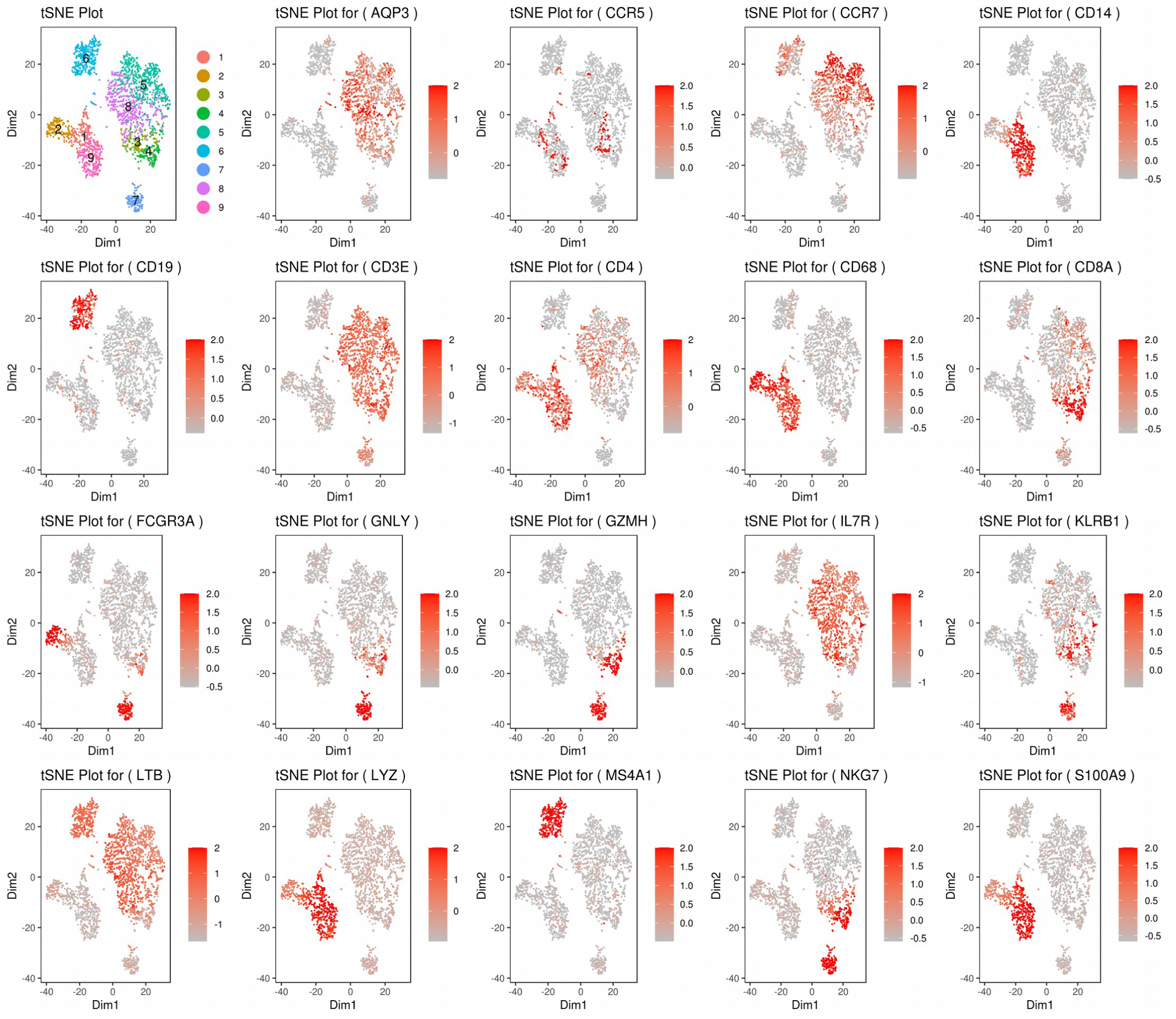
tSNE plots heatmapping the expression of some marker genes. As seen in the figure the cell communities do not have clear boundaries as seen in KNetL plots.

**Figure S2.**
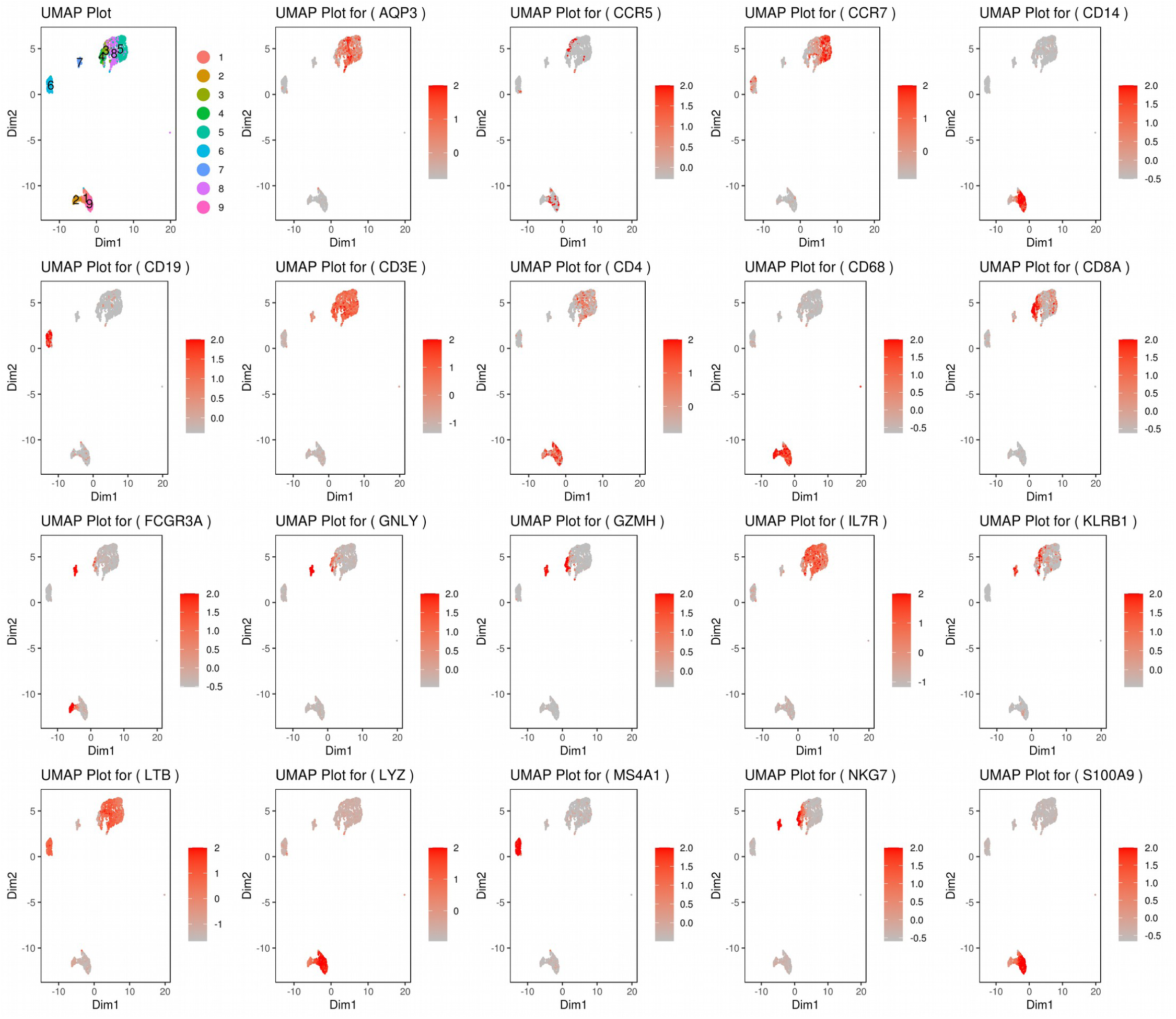
UMAP plots heatmapping the expression of some marker genes. As seen in the figure the cell communities do not have clear boundaries as seen in KNetL plots.

## Supplementary Codes

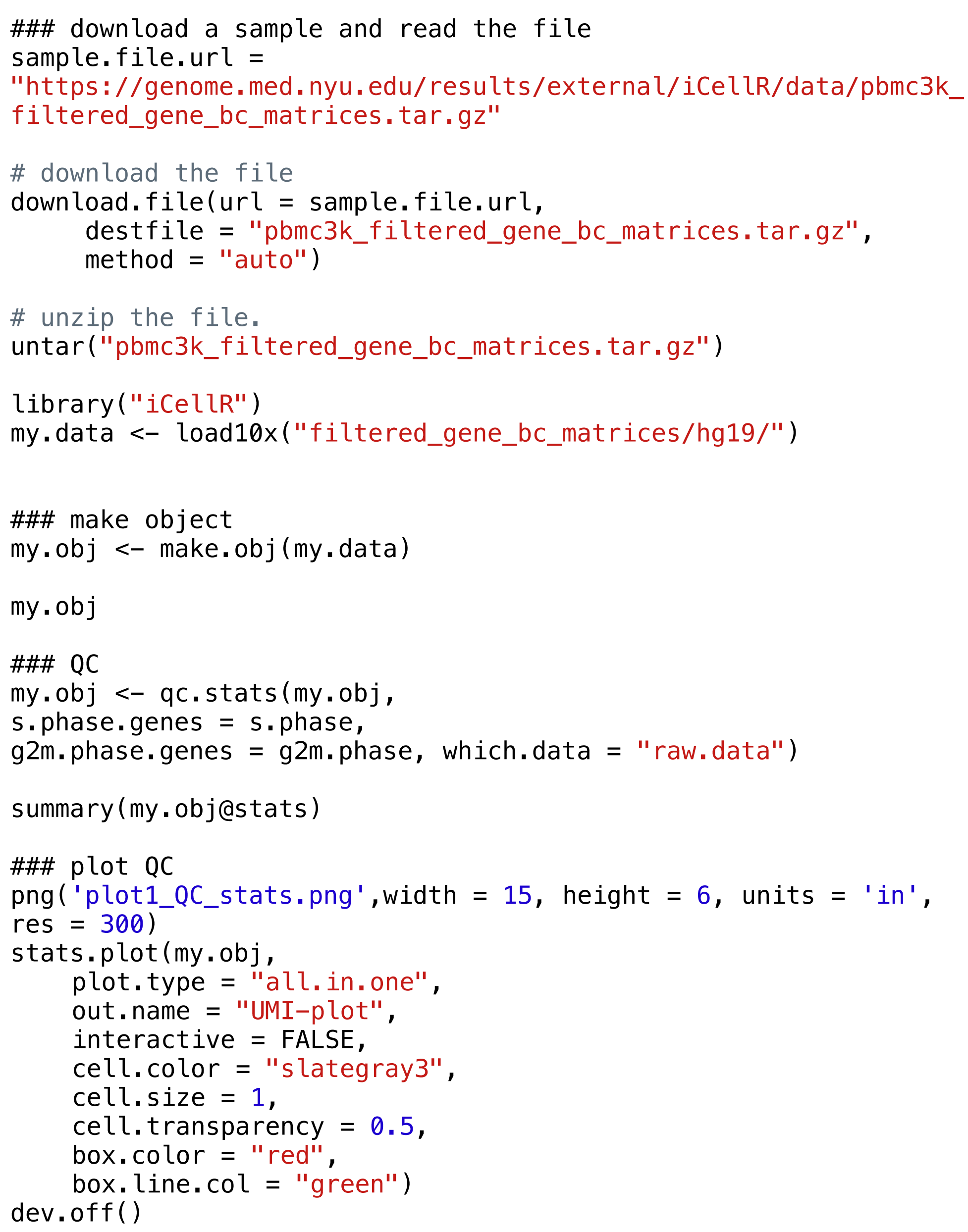

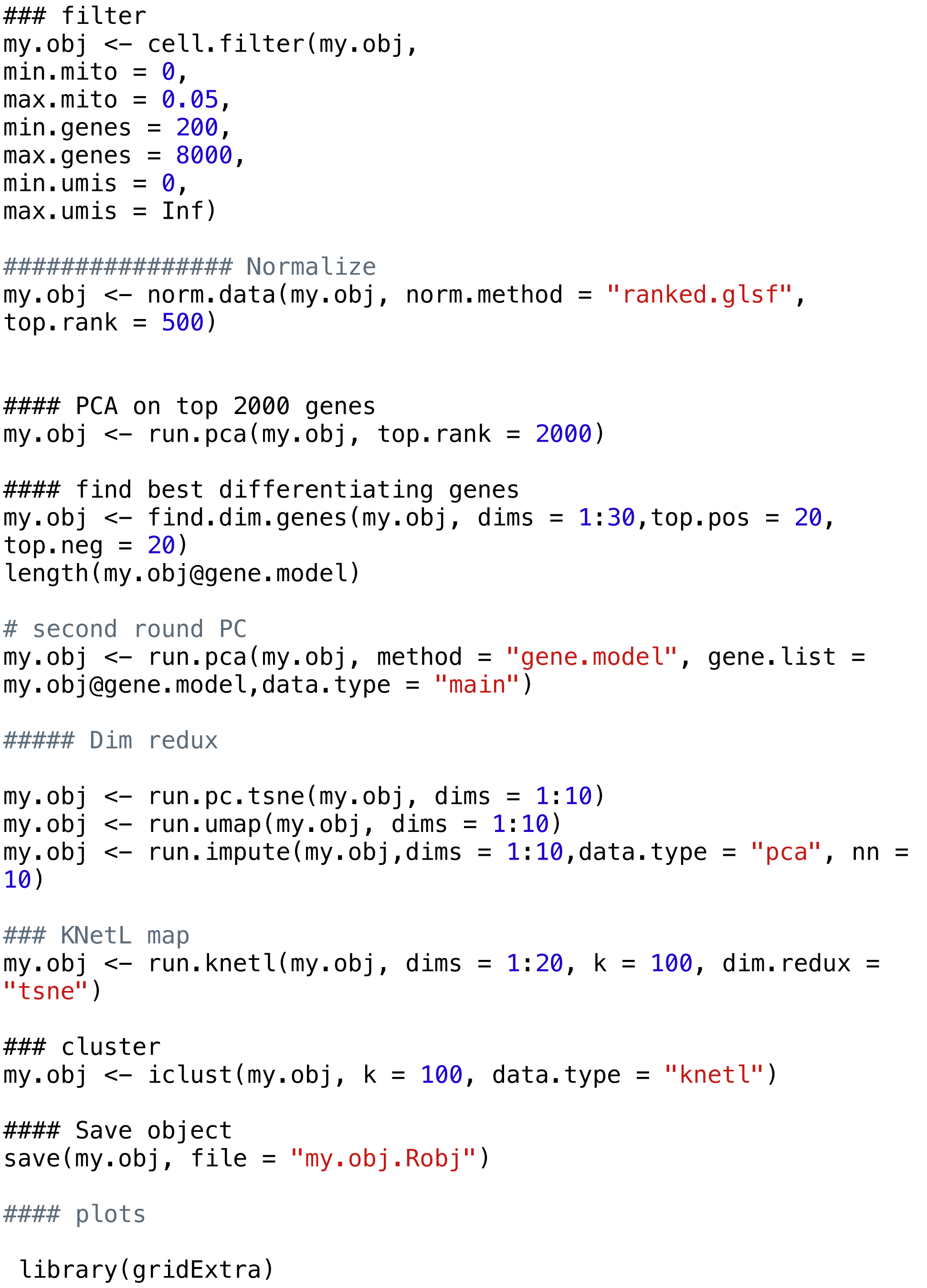

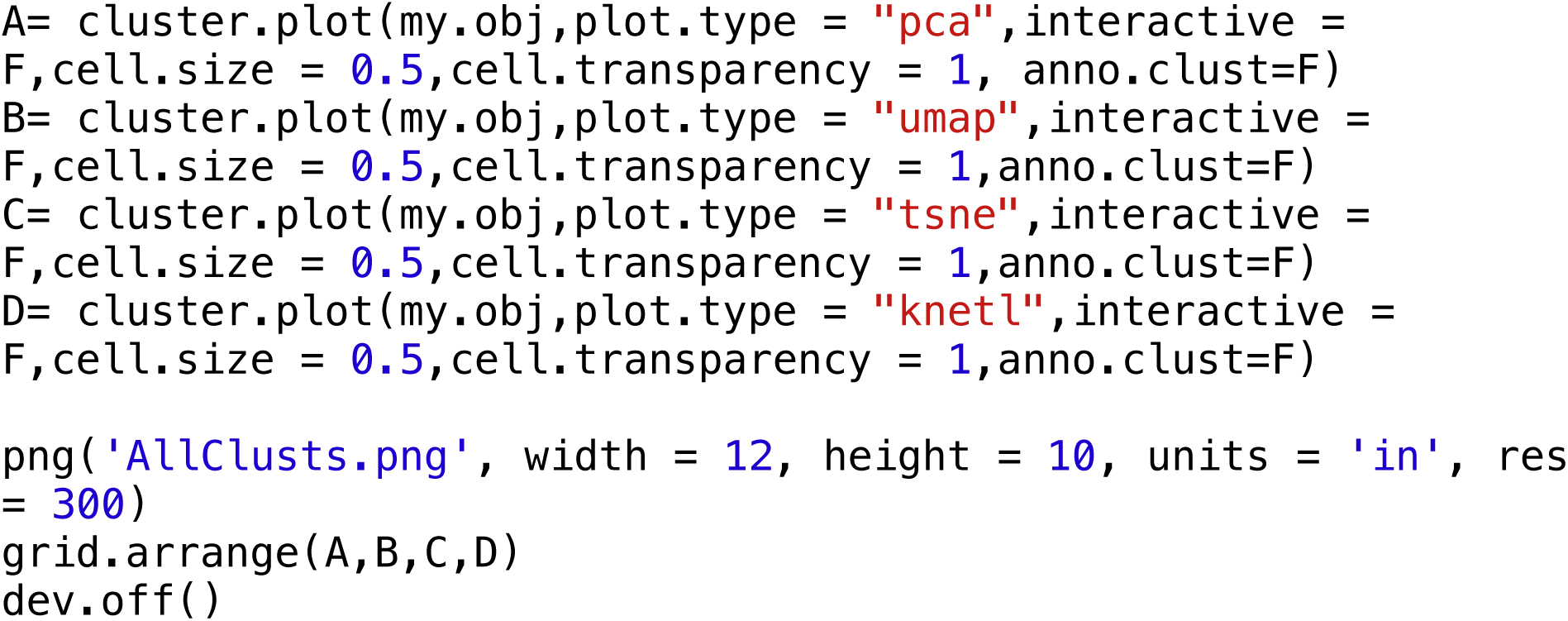

## REFERENCES

1- L.J.P. van der Maaten and G.E. Hinton. Visualizing High-Dimensional Data Using t-SNE. Journal of Machine Learning Research. 2008, 9(Nov):2579–2605.

2- Leland McInnes, John Healy, James Melville. UMAP: Uniform Manifold Approximation and Projection for Dimension Reduction. 2014, arXiv1802.03426.

3- Coifman RR, Lafon S, Lee AB, Maggioni M, Nadler B, Warner F, Zucker SW. Geometric diffusions as a tool for harmonic analysis and structure definition of data: diffusion maps. Proc Natl Acad Sci U S A. 2005 May 24;102(21):7426–31.

4- Ehsan Amid, Manfred K. Warmuth. TriMap: Large-scale Dimensionality Reduction Using Triplets. 2019, arXiv1910.00204.

5- Regev A, Teichmann SA, Lander ES, Amit I, Benoist C, Birney E, Bodenmiller B, Campbell P, Carninci P, Clatworthy M, Clevers H, Deplancke B, Dunham I, Eberwine J, Eils R, Enard W, Farmer A, Fugger L, Göttgens B, Hacohen N, Haniffa M, Hemberg M, Kim S, Klenerman P, Kriegstein A, Lein E, Linnarsson S, Lundberg E, Lundeberg J, Majumder P, Marioni JC, Merad M, Mhlanga M, Nawijn M, Netea M, Nolan G, Pe’er D, Phillipakis A, Ponting CP, Quake S, Reik W, Rozenblatt-Rosen O, Sanes J, Satija R, Schumacher TN, Shalek A, Shapiro E, Sharma P, Shin JW, Stegle O, Stratton M, Stubbington MJT, Theis FJ, Uhlen M, van Oudenaarden A, Wagner A, Watt F, Weissman J, Wold B, Xavier R, Yosef N; Human Cell Atlas Meeting Participants. The Human Cell Atlas. Elife. 2017 Dec 5;6. pii: e27041.

6- Khodadadi-Jamayran A, Pucella J, Zhou H, Doudican N, Carucci J, Heguy A, Reizis B, Tsirigos A. iCellR: Combined Coverage Correction and Principal Component Alignment for Batch Alignment in Single-Cell Sequencing Analysis. 2020, DOI: 10.1101/2020.03.31.019109

7- Fruchterman, T. M. J., & Reingold, E. M. Graph Drawing by Force-Directed Placement. Software: Practice and Experience, 1991, 21(11).

8- Jiarui Ding, Xian Adiconis, Sean K. Simmons, Monika S. Kowalczyk, Cynthia C. Hession, Nemanja D. Marjanovic, Travis K. Hughes, Marc H. Wadsworth, Tyler Burks, Lan T. Nguyen, John Y. H. Kwon, Boaz Barak, William Ge, Amanda J. Kedaigle, Shaina Carroll, Shuqiang Li, Nir Hacohen, Orit Rozenblatt-Rosen, Alex K. Shalek, Alexandra-Chloé Villani, Aviv Regev, View ORCID Profile Joshua Z. Levin. Systematic comparative analysis of single cell RNA-sequencing methods. 2019 Mar 23; doi: https://doi.org/10.1101/632216

9- Butler A, Hoffman P, Smibert P, Papalexi E, Satija R. Integrating single-cell transcriptomic data across different conditions, technologies, and species. Nat Biotechnol. 2018 Jun;36(5):411–420.

10- Haghverdi L, Lun ATL, Morgan MD, Marioni JC. Batch effects in single-cell RNA-sequencing data are corrected by matching mutual nearest neighbors. Nat Biotechnol. 2018 Jun;36(5):421–427.

